# Recognizing amino acid sidechains in a medium resolution cryo-electron density map

**DOI:** 10.1101/2024.12.10.627859

**Authors:** Dibyendu Mondal, Vipul Kumar, Tadej Satler, Rakesh Ramachandran, Daniel Saltzberg, Ilan Chemmama, Kala Bharath Pilla, Ignacia Echeverria, Benjamin M. Webb, Meghna Gupta, Klim Verba, Andrej Sali

**Affiliations:** Department of Bioengineering and Therapeutic Sciences, University of California, San Francisco, San Francisco, CA 94158, USA; Department of Cellular and Molecular Pharmacology, University of California, San Francisco, San Francisco, CA 94158, USA; Department of Pharmaceutical Chemistry, University of California, San Francisco, San Francisco, CA 94158, USA; Quantitative Biosciences Institute, University of California, San Francisco, San Francisco, CA 94158, USA; Department of Biochemistry and Biophysics, University of California, San Francisco, San Francisco, CA 94158, USA; Department of Synthetic Biology and Immunology, National Institute of Chemistry, 1000 Ljubljana, Slovenia

**Keywords:** cryo-electron microscopy, sequence threading, protein structure modeling, integrative structure modeling

## Abstract

Building an accurate atomic structure model of a protein into a cryo-electron microscopy (cryo-EM) map at worse than 3 Å resolution is difficult. To facilitate this task, we devised a method for assigning the amino acid residue sequence to the backbone fragments traced in an input cryo-EM map (*EMSequenceFinder*). *EMSequenceFinder* relies on a Bayesian scoring function for ranking 20 standard amino acid residue types at a given backbone position, based on the fit to a density map, map resolution, and secondary structure propensity. The fit to a density is quantified by a convolutional neural network that was trained on ∼5.56 million amino acid residue densities extracted from cryo-EM maps at 3-10 Å resolution and corresponding atomic structure models deposited in the Electron Microscopy Data Bank (EMDB). We benchmarked *EMSequenceFinder* by predicting the sequences of 58,044 distinct L-helix and β-strand fragments, given the fragment backbone coordinates fitted in their density maps. *EMSequenceFinder* identifies the correct sequence as the best-scoring sequence in 77.8% of these cases. We also assessed *EMSequenceFinder* on separate datasets of cryo-EM maps at resolutions from 4 to 6 L. The accuracy of *EMSequenceFinder* (63.5%) was better than that of three tested state-of-the-art methods, including *findMysequence* (45%), ModelAngelo (27%), and *sequence_from_map* in *Phenix* (12.9%). We further illustrate *EMSequenceFinder* by threading the SARS-CoV-2 NSP2 sequence into eight cryo-EM maps at resolutions from 3.7 to 7.0 Å. *EMSequenceFinder* is implemented in our open-source *Integrative Modeling Platform* (IMP) program. Thus, it is expected to be helpful for integrative structure modeling based on a cryo-EM map and other information, such as models of protein complex components and chemical crosslinks between them. *EMSequenceFinder* is available as part of our open source IMP distribution at https://integrativemodeling.org/.

## Introduction

Determining structures of biomolecular systems at atomic resolution is generally helpful in understanding their architectures, functions, and evolution as well as for modulating and designing their structures and functions. Cryo-electron microscopy (cryo-EM) has become a key method for structural characterization of large systems at near atomic resolution [1,2]. The process of *de novo* structure modeling based on a cryo-EM map generally proceeds in three steps [3,4]. First, backbone fragments are identified in the density map (step 1). Second, the amino acid residue sequence is assigned to the traced backbone fragments (step 2). Finally, loops that connect the traced backbone fragments and sidechains are added to obtain a complete and refined atomic model (step 3).

Building a protein structure model based on a cryo-EM map has been partially automated for higher resolution density maps (<3.5 Å) by traditional *de novo* modeling methods, such as *MAINMAST* [5], *Phenix* [6], and *Rosetta* [7], as well as deep-learning methods, such as *DeepTracer* [8] and *ModelAngelo* [9]. For 2-4 Å density maps, these methods typically produce models with 1-2 Å RMSD deviation from high-resolution atomic structures determined by X-ray crystallography, covering approximately 80-85% of the target sequence [9]. In addition, several automated methods have recently been developed and optimized for using high-resolution cryo-EM maps at better than 3.5 Å. For instance, *CryoDRGN* [10] employs deep generative networks to accurately reconstruct flexible regions in large macromolecular assemblies. Another approach, *Cryo2Struct* [11], utilizes a 3D transformer to predict atomic positions and amino acid residue types, followed by a Hidden Markov Model (HMM) protocol to build a backbone model. Another method, *DeepMainmast* [12], integrates *AlphaFold2* with a density tracing protocol to generate a complete atomic model. In parallel with advances in model building, various tools have been developed to validate the quality of structural models based on cryo-EM maps. For example, EMRinger [13] and DQScore [14] assess model accuracy by evaluating side-chain rotamers and atomic-level fit to the density map while also considering the principles of stereochemistry. In contrast, DAQ-score [15] applies a convolutional neural network (CNN) to evaluate local density quality around residues, providing a data-driven complement to traditional validation metrics.

Single particle cryo-EM reconstruction often yields maps in the medium resolution range of 4-8 Å [16], primarily due to the compositional and/or conformational heterogeneity of a cryo-EM sample; for instance, among the 37,178 cryo–EM maps in Electron Microscopy Database (EMDB) [17], 6,668 (17.9%), 22,448 (60.4%), and 8,062 (21.7%) are at <3 Å, 3-8 Å, and >8 Å resolution, respectively, as of September 2024. *De novo* modeling methods currently struggle to generate accurate and complete atomic models based on density maps with resolutions worse than 4 Å [18]. In the 4-8 Å resolution range, it is possible to trace most of the backbone using existing tools, such as Coot [19] and *Phenix* [6], usually with some manual intervention. However, existing methods often fail in assigning the sequence to the backbone, precluding computing an atomic model (step 3). Although these limitations can be mitigated with manual modeling, such interventions are generally time-consuming, not reproducible, and may fail to produce accurate and complete atomic models. Hence, there is a need for methods that can compute accurate and complete sequence threadings for the backbone fragments in medium resolution maps (step 2).

Identification of amino acid sidechains on a backbone fitted in a medium map resolution requires ranking alternative sidechains at a given position in a map. Tools, such as *Phenix* that employ the Resolve approach [20], compute the probability of each amino acid residue type at a given backbone position from modeling alternative sidechain rotamers in the map density using a Bayesian scoring function. While this direct approach is effective at map resolutions higher than 3.5 Å, where densities of most residues are well-resolved, it becomes less reliable at medium resolutions. At such resolutions, sidechain densities are often poorly resolved or absent, rendering direct methods less effective. Consequently, most sequence threading tools are limited to maps at resolutions better than 4 Å.

Unlike direct fitting methods, deep learning approaches leverage previously generated data to identify trends and features that may not be readily apparent [21]. CNNs are one such state-of-the-art method, widely used in image classification [22]. CNN-based methods, such as *DeepTracer* [8], *findMySequence* [23], and *ModelAngelo* [9], have already been applied to identify features in cryo-EM maps [24–26]. While these methods are promising, none were trained at map resolutions worse than 4 Å, potentially creating an opportunity for further method development.

Here, we constructed a Bayesian scoring function for ranking the 20 standard residue types based on the fit to a density map, map resolution, and secondary structure propensity. The density map fit was quantified by a CNN that was trained on ∼5.56 million amino acid residue densities extracted from cryo-EM maps and corresponding atomic structure models deposited in EMDB [27]. We used this Bayesian scoring function to solve the second (step 2) of the three steps in *de novo* structure modeling based on a cryo-EM map: the sequence is assigned to the traced backbone fragments by identifying the best scoring sequence threading among all possible threadings.

We found that our method (*EMSequenceFinder*) outperformed state-of-the-art methods in sequence threading when used with maps between 4 and 6 L resolution. *EMSequenceFinder* accurately assigned the sequence for 77.8% of backbone fragments (α-helices and β-strands). We further illustrated *EMSequenceFinder* by applying it to eight cryo-EM maps of the Severe Acute Respiratory Syndrome Coronavirus 2 (SARS-CoV-2) Non-Structural Protein 2 (NSP2) at resolutions between 3.7 and 7.0 L. The assignment accuracy of *EMSequenceFinder* ranged from ∼95% at the resolution of 4 L to ∼50% at the resolution of 6 L, again demonstrating superior performance compared to other tested methods.

## Methods

We developed and benchmarked *EMSequenceFinder*, a method for assigning protein sequences to backbone fragments fitted into a density map (backbone traces) (**Fig. 1**). Specifically, the input consists of one or more protein sequences, their cryo-EM density map with a resolution better than approximately 8 Å, and backbone traces for α-helices and β-strands. The output consists of the best sequence threadings for each backbone fragment (step 2). This output is generated by ranking alternative threadings of the sequences to the backbone traces, partly based on scoring how well sidechains fit the map using a CNN. A flowchart of *EMSequenceFinder* as implemented in our open-source *Integrative Modeling Platform* (IMP) software (https://integrativemodeling.org) [28–30] is shown in **Figure 1**.

**Figure 1.**
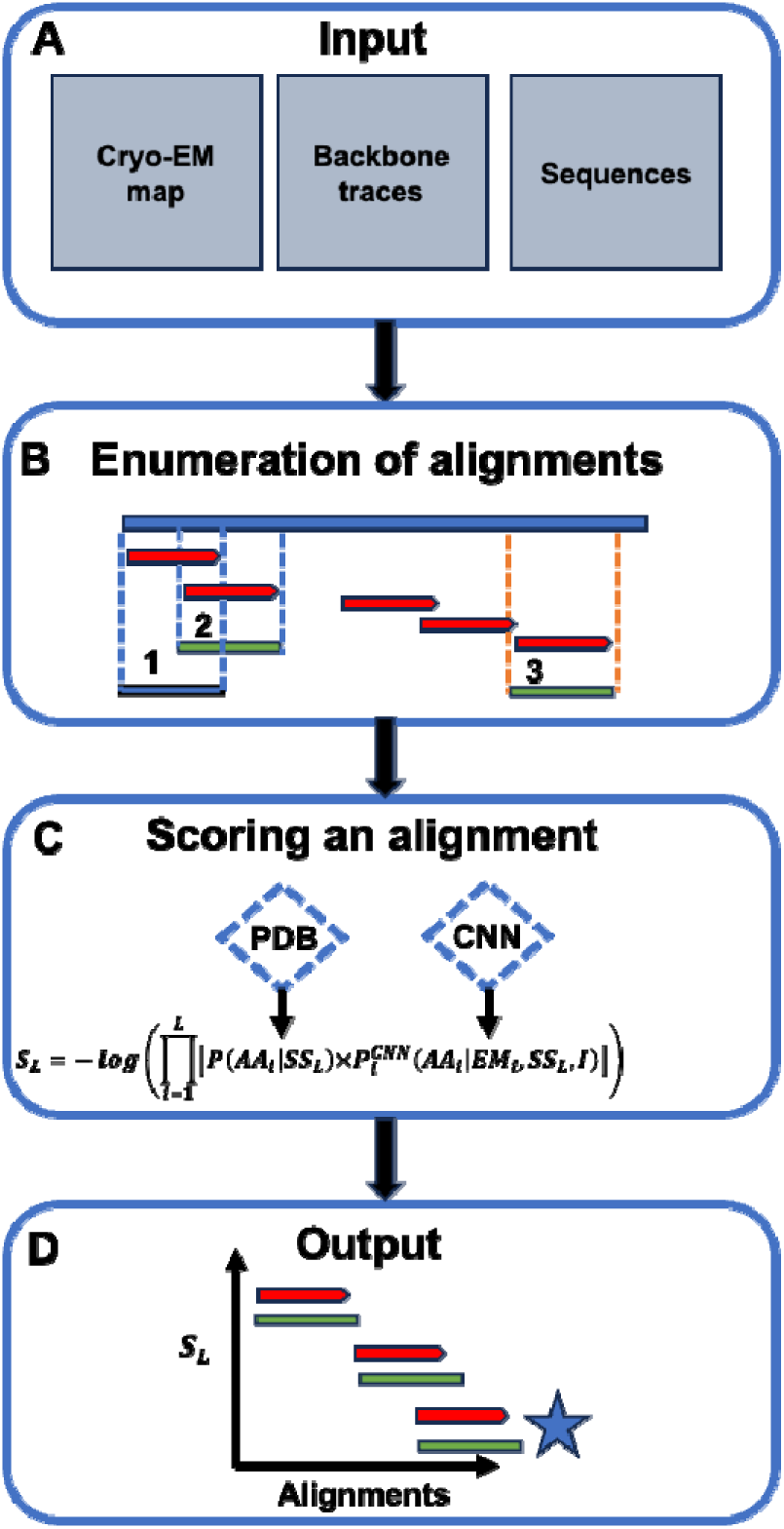
Flowchart of *EMSequenceFinder*. **(A**) The input consists of a cryo-EM map, backbone traces (L-helices and/or β-strands fit into the cryo-EM map), and protein sequence(s). (**B**) For a given backbone trace, all gapless threadings (in red bars) of the input protein sequences are enumerated. (**C**) Each sequence-backbone trace threading (alignment) is scored using a Bayesian scoring function that consists of two terms, including secondary structure propensity of each residue in the sequence () and a fit of each residue to the map as evaluated by a CNN (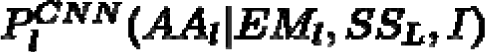. (**D**) The output is the best-scoring threading for each input backbone trace.

### Input data

The input consists of three items. The first input is one or more protein sequences to be localized in a cryo-EM map, with no limitations on the number or length of sequences other than those imposed by computer memory and time. Typically, *EMSequenceFinder* can localize a protein with 1,000 residues in under 100 seconds of CPU time.

The second input is a cryo-EM map in the standard MRC format at resolution better than 8 Å. While there is no limit on the best resolution, aligning a sequence to backbone traces in maps at better than 2.5 - 3.0 Å resolution is straightforward, without the need for generating and scoring many alternative sequence threadings, as performed in *EMSequenceFinder*.

The third input is a set of α-helical and β-strand backbone fragments either traced or fitted into the map (backbone traces). These backbone traces should specify the positions of the corresponding amide N, C___, C___, as well as carbonyl C and O atoms. They can be obtained manually or by using programs, such as *Phenix* [6], *Rosetta* [7], *DeepTracer* [8], *SSEHUNTER* and *SSEBUILDER* [31], *ModelAngelo* [9], and *Cryo2Struct* [11]. In this work, we specifically trained our CNN models using density maps for α-helical and β-strand backbone fragments, as obtaining reliable maps for other secondary structure elements, such as coils and loops, remains more challenging. Coil/loop regions, especially in medium to low-resolution maps, often do not fit the maps well. In the future, irregular fragments could be modeled in step 3, as mentioned in **Introduction**.

### Enumeration of threading

Each backbone trace is considered independently from other traces. For a given trace, all gapless threadings of input protein sequences are enumerated. Specifically, the trace is aligned with all possible contiguous segments of each input sequence, without gaps. Thus, for an input sequence of *N* residues, *EMSequenceFinder* generates (*N - L* + 1) alternative threadings through the trace, where *L* is the number of residue positions in the trace. Each alternative threading is scored as described below and ranked by the score. If the correct sequence ranked within the top *L*/2 scores, we deemed the prediction accurate. Our threading procedure was applied one fragment at a time, independently for each fragment. When applied to all fragments in a protein, the accuracy of our approach was evaluated by counting the number of fragments correctly assigned to their respective positions.

### Scoring a threading

The score for a given backbone trace - sequence threading is:

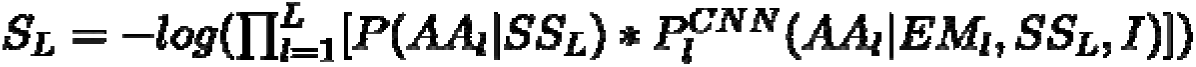

where the product runs over all ***L*** residue positions (***l***) in the trace; ***AA_l_*** is the type of the residue at trace position ***l***; ***EM_l_*** is the map density at trace position ***l***; ***SS_l_*** is the secondary structure type of the backbone trace; and ***I*** is other information. In the current implementation, we only used map resolution as an additional parameter. Although we also experimented with incorporating local resolution calculated with Phenix, along with the overall map resolution, we did not observe a significant improvement in prediction accuracy. As a result, we did not include local resolution information into the scoring function.

At each trace position, we multiply two terms. The first term, ***P(AA_l_|SS_L_***), is the probability of the aligned residue type, given ***SS_L_*** (these probabilities for the 20 standard residue types sum up to 1). This first term is extracted from a table, described below.

The second term, 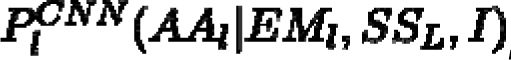, is the probability of the aligned residue type, given ***EM_l_, SS_L_***, and ***I*** (these probabilities for the 20 standard residue types sum up to 1). This second term is computed by our CNN model, described below.

### Residue type propensity for each secondary structure designation

The probability of occurrence of each of the 20 standard residue types in each of the three secondary structure designations was estimated from a statistical sample that included all assigned residues in all protein chains in the Protein Data Bank (PDB) [32] in early 2020:

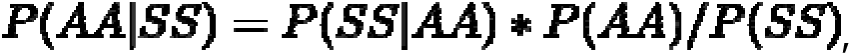

where ***P***(***AA***) is the fraction of each of the 20 standard residue types in a statistical sample of residues of defined type and secondary structure type, ***P***(***SS***) is the fraction of residues for each secondary structure type, and ***P***(***SS|AA***) is the fraction of secondary structure types for each residue type. Secondary structure types (H for helix, E for strand, and C for coil) were computed by DSSP [33].

### Convolutional neural network for residue type prediction

A CNN that computes the likelihood of each residue type, given the sidechain voxel intensities and map resolution, was trained using the Keras [34] API of the TensorFlow software package [35]. The neural network architecture consists of four blocks (**Fig. 2**). The input is the sidechain density of a residue (***l***), represented as a 14×14×14 matrix of values of normalized voxel intensities (***EM_l_***). The final output layer with softmax activation output is a vector with 20 elements representing the likelihood of each amino acid residue type (***AA_l_***) as 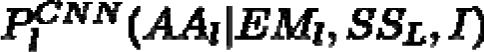, where is additional information (here, only the map resolution).

**Figure 2.**
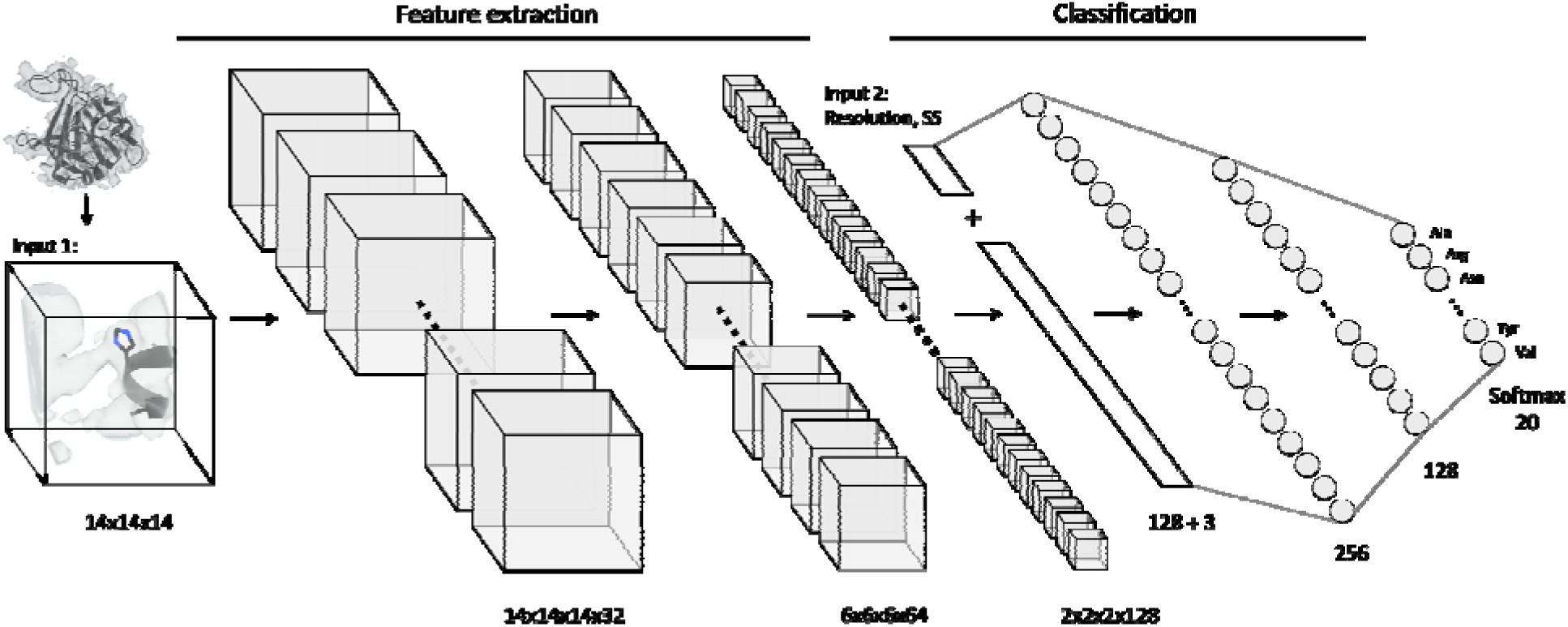
The Convolutional Neural Network (CNN) architecture used in EMsequenceFinder. The 3D CNN architecture consists of 4 blocks. The first 3 blocks were used to extract features from the residue sidechain densities, whereas the last block was used to predict probabilities of residue types from a concatenated input of the extracted features from the first 3 blocks and other information (map resolution and secondary structure type). Each of the first three blocks consist of two convolutional layers followed by batch normalization and a dropout layer. The input for these blocks is the sidechain density of a residue (***l***), represented as a 14×14×14 matrix of values of normalized voxel intensities (***EM_l_***). The three blocks are used to learn features from the density information. The output vector from a dense layer of the third convolution block is fed to a global average pooling layer. The output from the dense layer is then concatenated with additional input data, including resolution of the map and secondary structure type (***SS_L_***) of the trace containing residue and fed to the fourth block. The fourth block consists of two fully connected dense layers. The final output layer with softmax activation outputs a set of 20-dimensional vectors representing the likelihood of each amino acid residue type (***AA_l_***) as 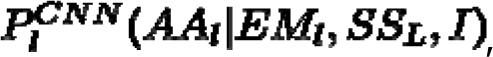, where, ***I*** is other information (here, the map resolution).

Categorical cross-entropy (CCE), which minimizes the average cross-entropy across the entire training dataset, was used as the loss function for training the neural network. This loss function incentivizes higher probabilities for the target class, regardless of whether they are the maximum probability. A learning rate of 0.00008 for changing the network weights was chosen as a compromise between the stability and convergence time of training. The Adam optimizer [36] was used to iteratively update the network weights. A batch size of 64 was used to allow for calculation on our 24GB Titan GPU hardware. We chose a maximum of 100 epochs of training, with an early stopping criterion of 10 epochs if there was no improvement.

The distribution of amino acid residue types for each secondary structure type is uneven. Thus, we weighted the score of each residue type by the inverse of its count in the training set. To guard against overfitting, we set aside 10% of our database for validation (validation set), ensuring that the threading accuracy and CCE were not significantly worse than those for the training set (80% of the database). The remaining 10% of the database was used for testing (Test Set I).

### Database for training, validating, and testing the convolutional neural network

A database of sidechain densities determined by fitting molecular models into cryo-EM maps was prepared to train, validate, and test our CNN model. Each entry consisted of voxel intensities for a 14 Å x 14 Å x 14 Å cube centered on the C_β_ atom (**Fig. 2**), sampled on a 1 Å grid, for a total of 2,744 voxels. Each sidechain entry also contained additional information, including EMDB ID of the map, residue type, author-defined resolution of the entire map (Å), and secondary structure type. The database was constructed as follows.

We started with the EM maps and associated atomic models deposited in EMDB [17,35] as of 3/13/2021. We excluded EMDB entries with maps at worse than 10.0 Å and models with nucleic acids. We also excluded all poly-Ala segments and undefined sidechains. Each cryo-EM map was scaled to a common reference using histogram matching [37]. Map resolutions in our database ranged from 1.15 to 10 L. Next, backbone fragments were extracted from the atomic structure models. A backbone fragment is defined as a segment of at least 4 consecutive residues in the same secondary structure type, as defined by *STRIDE* [38]. Backbone fragments whose EMDB sequence did not match their UniProt sequences were excluded. The resulting dataset included 622,884 secondary structure elements and 5,542,632 residues from 3,914 EMDB entries (out of 5,515 total entries with sample type “Protein” in EMDB); 4,188,243 and 1,354,390 residues are in L-helices and β-strands, respectively (**Fig. 3**).

**Figure 3.**
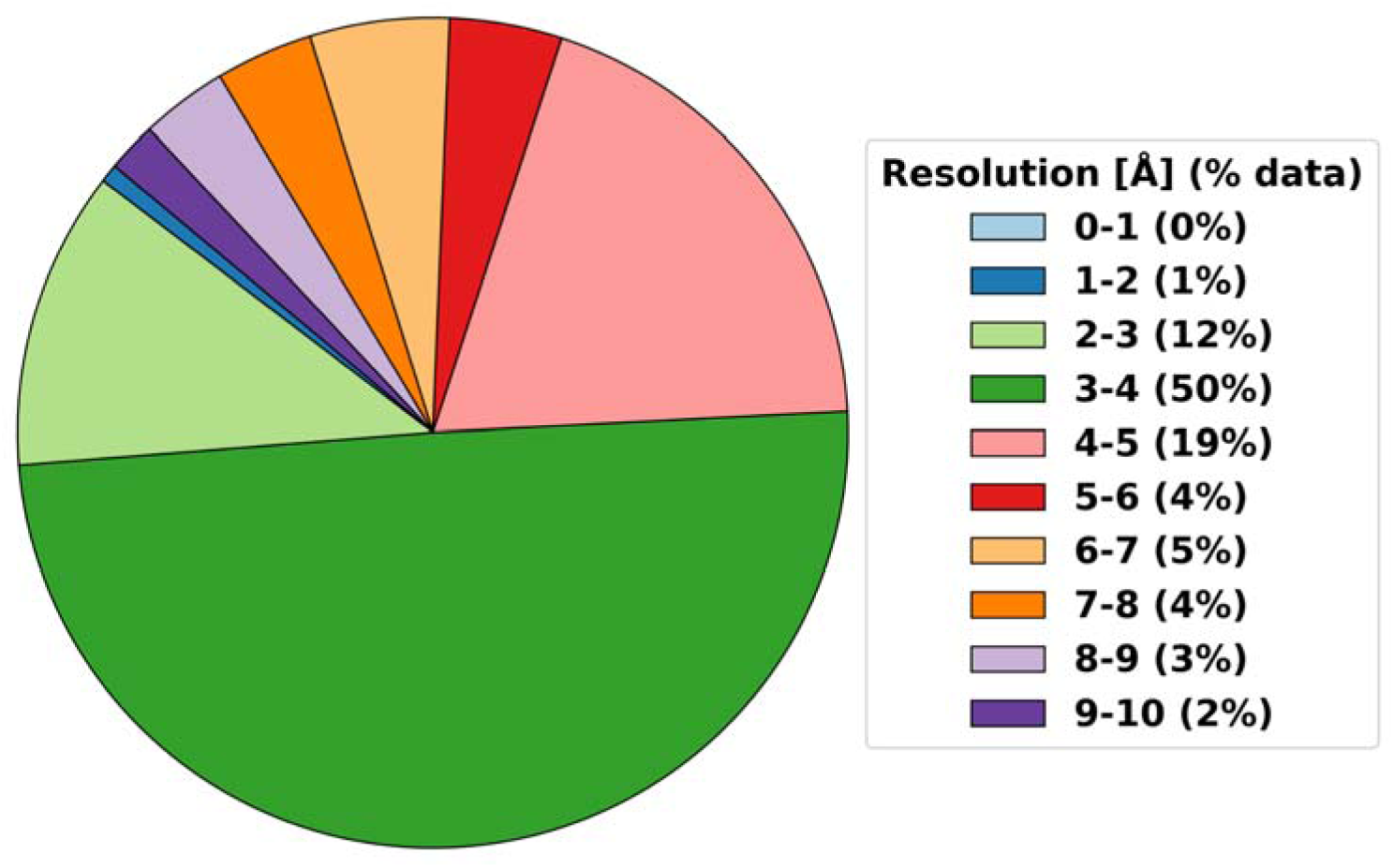
The number of residues extracted from 3914 EMDB entries used to train the CNN model after processing. Most maps have resolutions from 3 to 4 L. Only 18% of the maps are at resolutions worse than 5 L. Percentage of L-helical and β-strand residues in the corresponding EMDB atomic structures are 75.6% and 24.4%, respectively.

We then extracted the density in the 14 Å x 14 Å x 14 Å cube around each residue in each backbone fragment from the corresponding map. For Gly residues, the virtual C_β_ atom position was computed from the positions of the CA, C, and N atoms. The scaled cryo-EM map was linearly interpolated to obtain the densities at the voxels in the 14 Å x 14 Å x 14 Å cube, stored in the database as a vector with 2,744 elements.

Finally, we constructed a second test set (Test Set II) using cryo-EM maps and atomic models with an EMDB release date between 8/14/2021 and 3/14/2023, resolution between 3 to 8 Å, and a molecular weight of up to 250 kDa. This test set contains 742,780 residues in 58,044 distinct fragments from 1,251 cryo-EM maps.

### Databases for comparing *EMSequenceFinder* with state-of-the-art methods

To compare *EMSequenceFinder* with state-of-the-art methods, we prepared two datasets of cryo-EM maps: one with resolutions between 4.0 and 4.5 Å (Test Set IIIa) and another one with resolutions between 5.0 and 6.0 Å (Test Set IIIb) (Supplementary Table S1). Test Set IIIa was created by selecting all maps with resolution between 4.0 and 4.5 Å from the dataset described in [39]. This study evaluated the ability of *Phenix,* one of the state-of-the-art methods considered here, to automatically build an atomic model based on a cryo-EM map at a resolution from 2 to 4.5 Å. To further assess performance on maps with resolutions worse than 4.5 Å, we compiled Test Set IIIb from maps deposited between August 2021 and September 2022, with resolutions ranging from 5.0 to 6.0 Å and protein molecular weights of less than 25 kDa. We set the upper limit to 6.0 Å to explore a potential strength of *EMSequenceFinder* in comparison to the state-of-the-art methods tested here, which were recommended only for maps at resolutions of better than 4.5 Å. In the current study, we did not thread multiple copies of the same protein sequence. However, our tracing algorithm is easily extensible to systems with multiple copies of the same sequence.

#### Model output

The output of *EMSequenceFinder* for each secondary structure fragment is the best ranked (lowest scoring) threading of the trace, provided as a text file. Additionally, a 3D coordinate file containing sidechain coordinates can be computed using IMP.

## Results and Discussion

We first assess a key aspect of *EMSequenceFinder*, namely its ability to predict the type of the residue, given the sidechain cryo-EM density, probable secondary structure type near that density, and the resolution of the map. We then follow up by assessing the ability of *EMSequenceFinder* to predict the sequence of a single fragment. We conclude the assessment by quantifying the performance of *EMSequenceFinder* in assigning one or more protein sequences to backbone fragments fitted into a density map of the entire system. Finally, we demonstrate the utility of *EMSequenceFinder* by its application to a real-world case of modeling the SARS-CoV-2 Nsp2 protein based on cryo-EM maps at resolutions ranging from 3.7 to 7.0 Å.

### Accuracy of predicting type of a residue

We evaluated the performance of *EMSequenceFinder* in predicting individual amino acid residue types based on sidechain density using Test Set II. Importantly, Test set II does not overlap with the training validation datasets and Test Set I, which were used to develop the CNN model. *EMSequenceFinder* correctly identified 41% of the residues in Test Set II as the best-scoring residue type, a result comparable to the ∼47% accuracy achieved with Test Set I. This consistency indicates that the training dataset for the CNN model was sufficient and that the CNN model is robust.

Next, we quantified the performance of *EMSequenceFinder* by its Receiver Operating Characteristic (ROC) curve (**Fig. 4A**), which plots the true positive rate (TPR) on the Y-axis and false positive rate (FPR) on the X-axis. The larger the Area Under the Curve (AUC), the better the classifier. Our CNN model was trained for multiclass classification. Thus, we first binarized the output. In general, binarizing can be done in two different ways. First, the One-vs-Rest scheme compares each class (i.e., a specific amino acid residue type) against all the other classes, which are combined and treated as a single group. Second, the One-vs-One scheme compares every unique pairwise combination of classes [40]. Here, we report the results of both micro-averaging (data from all classes are averaged, for the One-vs-Rest scheme) and macro-averaging (metrics for each class are calculated independently and subsequently averaged) [41]. The AUC for both micro- and macro-averaging is notably high at 0.89 (**Fig. 4A**).

**Figure 4.**
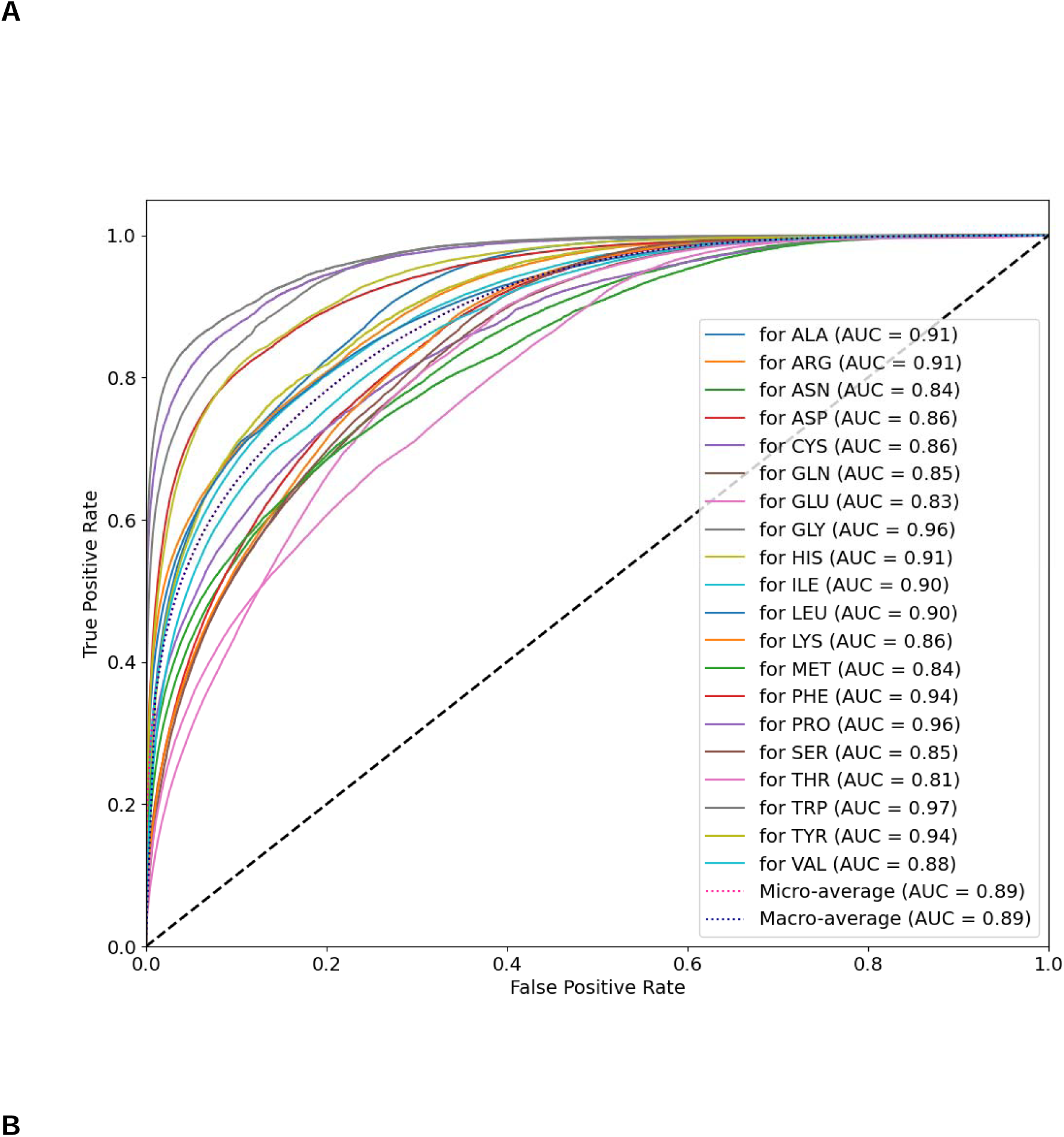
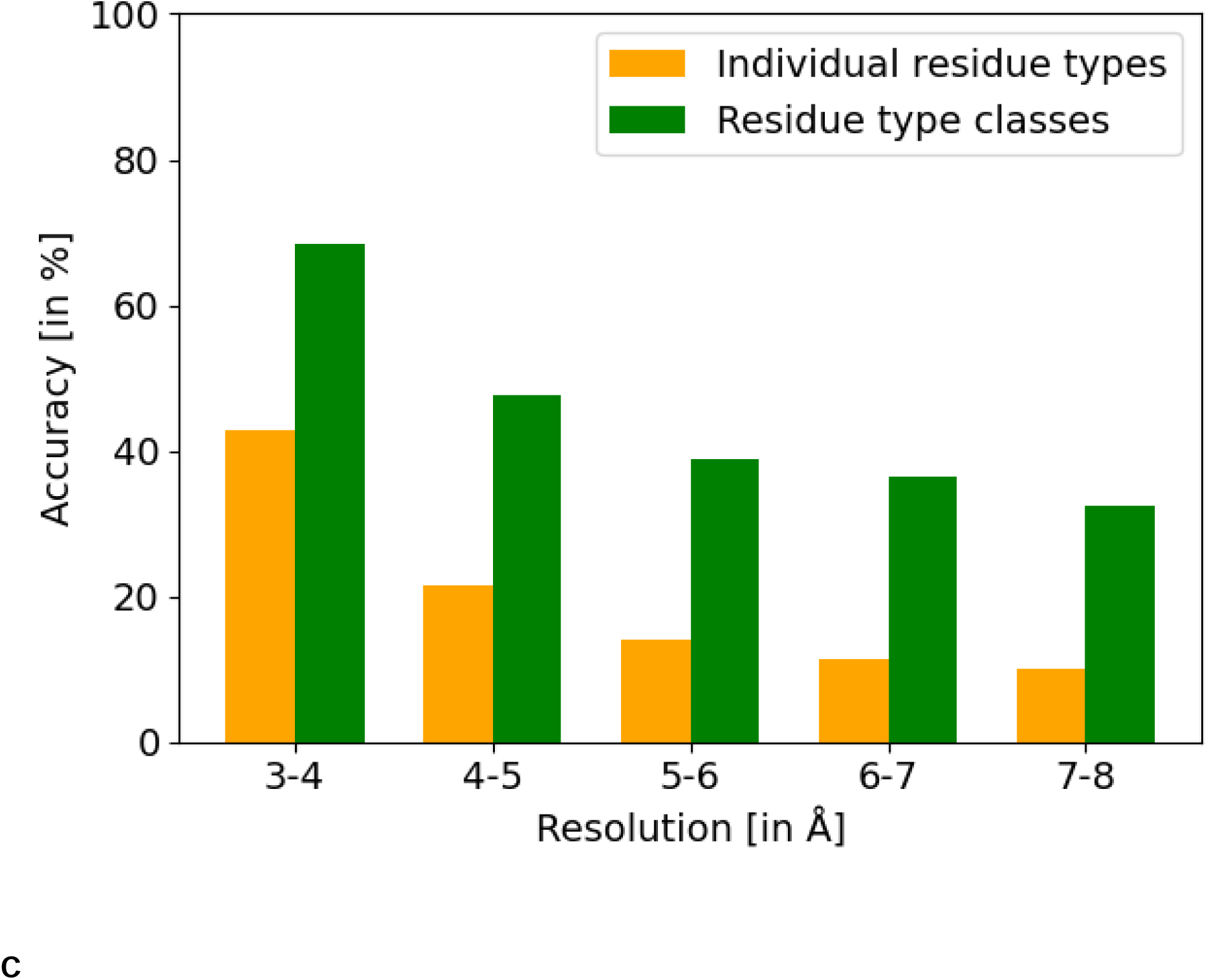
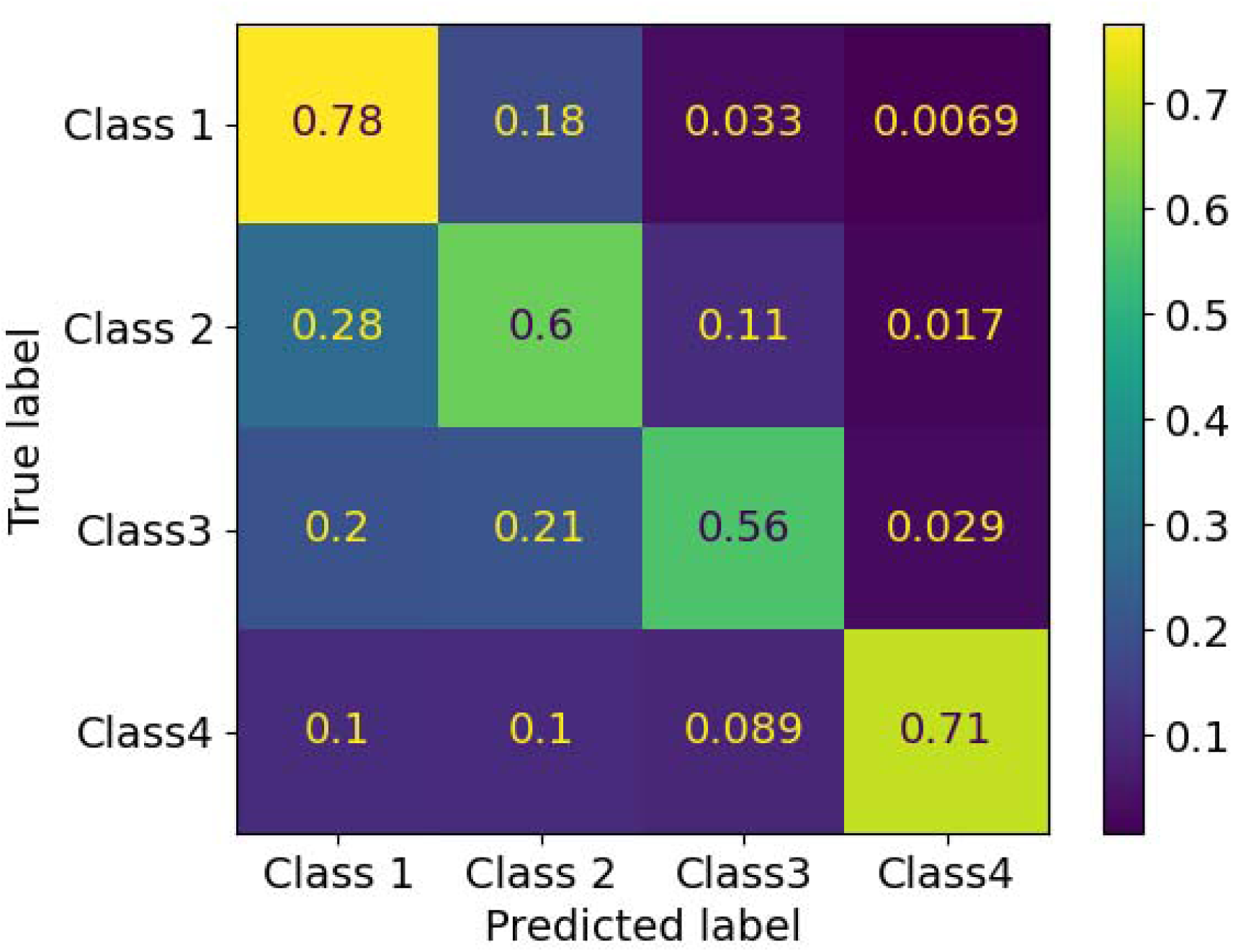
Assessment of amino acid residue type prediction accuracy of the CNN model. **(A)** The ROC curves show residue classification accuracy of the model for each residue type. **(B)** Prediction accuracy of the 20 standard residue types (orange) and residue type classes based on size (green; see text). (**C**) The confusion matrix for classification of residue types into the sized-based groups (the matrix is normalized over the true conditions (rows)).

We further evaluated the classifier performance as a function of map resolution, anticipating a decrease in prediction accuracy with a decrease in map resolution. Lower-resolution maps are less informative about the sequence due to higher fitting error in the backbone models; such maps are also underrepresented in our training set (**Fig. 3**). These factors negatively impact learning and generalization by the deep learning models. Indeed, we observed a gradual decrease in accuracy from 42% at 3-4 Å to 10% at 7-8 Å (orange bars in **Fig. 4B**). Thus, better modeling of noise in the input density map may improve prediction accuracy. Correspondingly, density maps produced by different methods, such as single particle reconstruction, subtomogram averaging, and 3D electron diffraction, may result in different prediction accuracies even at the same nominal resolution, due to different forms of noise. It is also conceivable that map processing methods, such as denoising and density modification [42], improve prediction accuracy, because they reduce noise.

Discriminating between sidechain types of similar shapes (*eg*, glutamine and asparagine) is particularly challenging, especially as the map resolution worsens. To characterize this challenge, we calculated the accuracy of predicting amino acid residue classes (based on their size) instead of the individual 20 standard residue types. The amino acid residue types were grouped with respect to their size to define four classes [24,43] (**Table 1**), as follows. Class 1 includes small and hydrophobic residues, such as glycine and alanine; Class 2 includes acidic and neutral polar residues, including aspartic acid and glutamine; Class 3 contains the basic residues arginine and lysine; and Class 4 consists of aromatic residues, such as histidine and tryptophan. Prediction accuracy for residue classes is significantly higher than that for individual residue types (green bars in **Fig. 4B)**. Similarly to the individual residue types, the prediction accuracy of residue classes also declines as the map resolution worsens. A confusion matrix demonstrates that basic amino acid residue types (Class 3) are the most misclassified residues, often misclassified with other acidic or branched amino acid residues (Class 2) (**Fig. 4C**).

**Table 1.** Grouping of amino acid residue types by the similarity of their sidechain map densities.

| Residue Type Class | Amino Acid Residue Types |
| --- | --- |
| Class1 | GLY, ALA, SER, CYS, VAL, THR, ILE, PRO |
| Class2 | LEU, ASP, ASN, GLU, GLN, MET |
| Class3 | LYS, ARG |
| Class4 | HIS, PHE, TYR, TRP |

### Accuracy of predicting the sequence of a fragment

We next assess *EMSequenceFinder* by benchmarking its accuracy in identifying the correct fragment sequence from all possible alternative fragment sequences, given the sidechain densities of residues in the fragment, residue secondary structure types, and the resolution of the map. Threading the correct sequence onto a fragment is anticipated to yield more accurate results than predicting individual residue types, because of a cumulative and synergistic effect of combining scores for multiple single residues constrained by sequence.

To perform the benchmark, *EMSequenceFinder* predicted probabilities for each of the 20 amino acid residue types at each residue position of each of the 58,044 distinct α-helices and β-strands in Test Set II, resulting in an *L* x 20 matrix for a fragment of *L* residues. This matrix is then used to score and rank all possible alternate threadings for the fragment (**Methods**).

The overall prediction accuracy for all fragments in Test Set II was 77.8%. Specifically, accuracies were 79.5% for the α-helix fragments and 75.7% for the β-strand fragments. As expected, prediction accuracy declines significantly with the resolution of the map (**Fig. 5A**). *EMSequenceFinder* performs reasonably well for maps with resolutions better than 6 Å. Interestingly, the decrease in accuracy was more pronounced for β-strands than α-helices. Possible reasons include a worse local map quality and larger backbone fitting errors for β-strands than α-helices, at the same overall map resolution.

**Figure 5.**
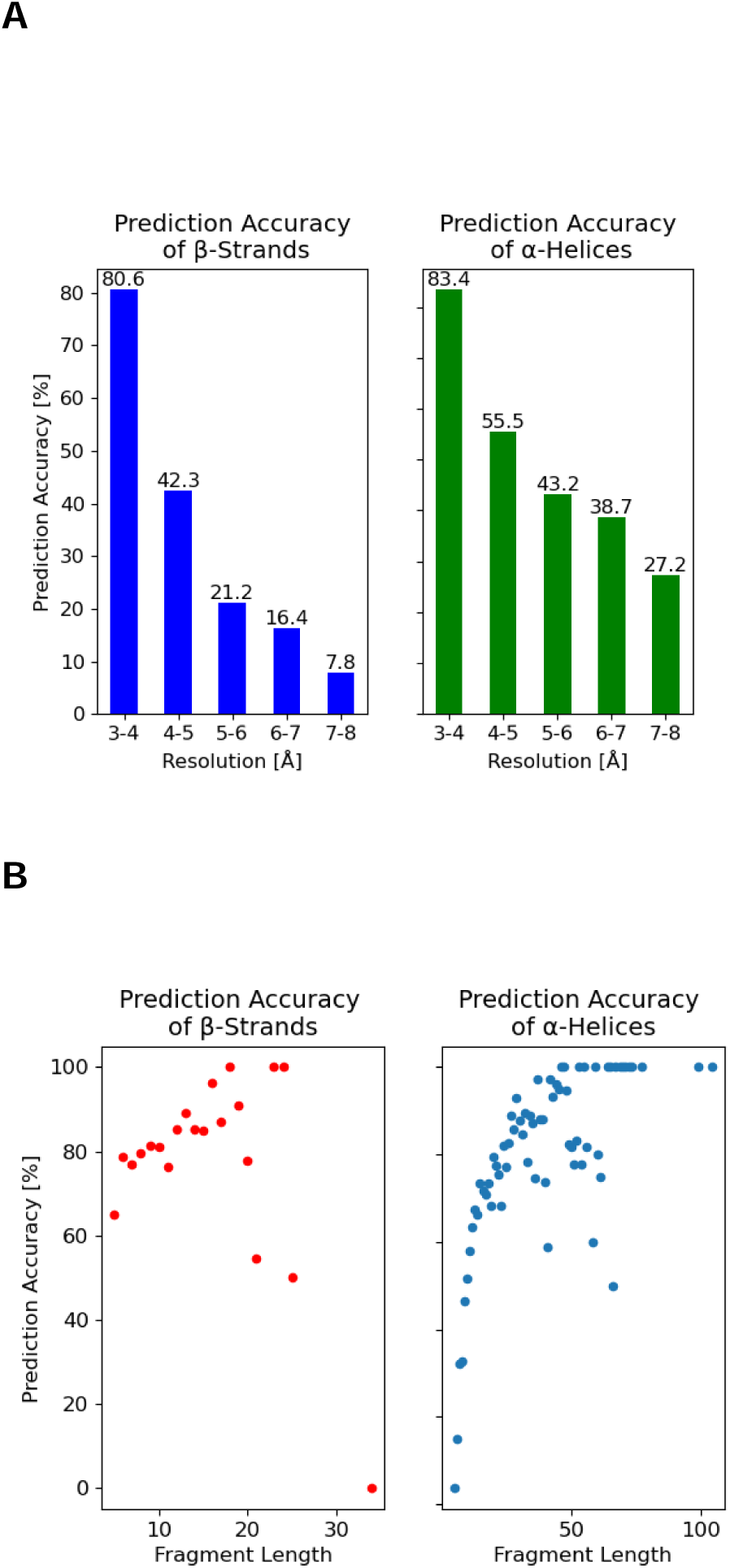
Mean prediction accuracy of the fragment sequences. **(A)** Mean prediction accuracy for L-helices (green) and β-strands (blue) for different map resolutions. **(B)** Mean prediction accuracy for L-helices (green) and β-strands (blue) as a function of fragment length.

We also examined the correlation between fragment length and prediction accuracy. α-helix fragments with at least 7 residues and β-strand fragments with at least 15 residues were predicted accurately in more than 60% of cases (**Fig. 5B**). For longer fragments, prediction accuracy approaches near perfection, as expected (above). Thus, for assigning the sequences of all input fragments, we propose an iterative threading approach that starts with assigning the sequence of the longest unassigned fragment, eliminates the assigned sequence from further consideration, and iteratively proceeds with the remaining fragments.

### Comparison of *EMSequenceFinder* with state-of-the-art methods

With a benchmark of *EMSequenceFinder* in predicting both individual amino acid residue types and fragment sequences in hand, we next compare it with state-of-the-art methods, including *findMySequence* [23], *ModelAngelo* [9], and *Phenix* [6].

*findMySequence* method was trained on maps with resolutions better than 4.5 Å, in contrast to *EMSequenceFinder* training range of 3-10 Å (**Introduction**). For assessment, we utilized two datasets: Test Set IIIa and Test Set IIIb (**Methods**). We used the *chain_comparison* program from *Phenix* [6], with default parameter settings, to calculate the percentage of the input sequence predicted correctly by each of the three tested methods. The backbone coordinates were obtained from the corresponding atomic models and were used to extract the sidechain densities. For this assessment, we considered only α-helices and β-strands, as identified by *STRIDE* [38]. The *ModelAngelo* models (using the build method with the input map and FASTA sequence) were directly compared to the fitted PDB models for the respective test set maps, using o*nly_keep_best_chain=True* in the chain_comparison tool of *Phenix*. Deep learning methods, *findMySequence, ModelAngelo,* and *EMSequenceFinder*, are significantly more accurate than *Phenix* in the 4-6 Å map resolution range (**Fig. 6**; **Supplementary Fig. S2**). *EMSequenceFinder* achieved an overall accuracy of 63.5%, while *findMySequence* and *ModelAngelo* accuracies were 45% and 27%, respectively. In contrast, the accuracy of the *sequence_from_map* function of *Phenix* was only 12.9%. *EMSequenceFinder*, f*indMySequence*, and *ModelAngelo* exhibited comparable accuracies on maps within the 4.0–4.5 Å resolution range. However, for maps with resolutions worse than 5 Å, *EMSequenceFinder* is more accurate than *findMySequence* and *ModelAngelo* (**Fig. 6B)**.

**Figure 6.**
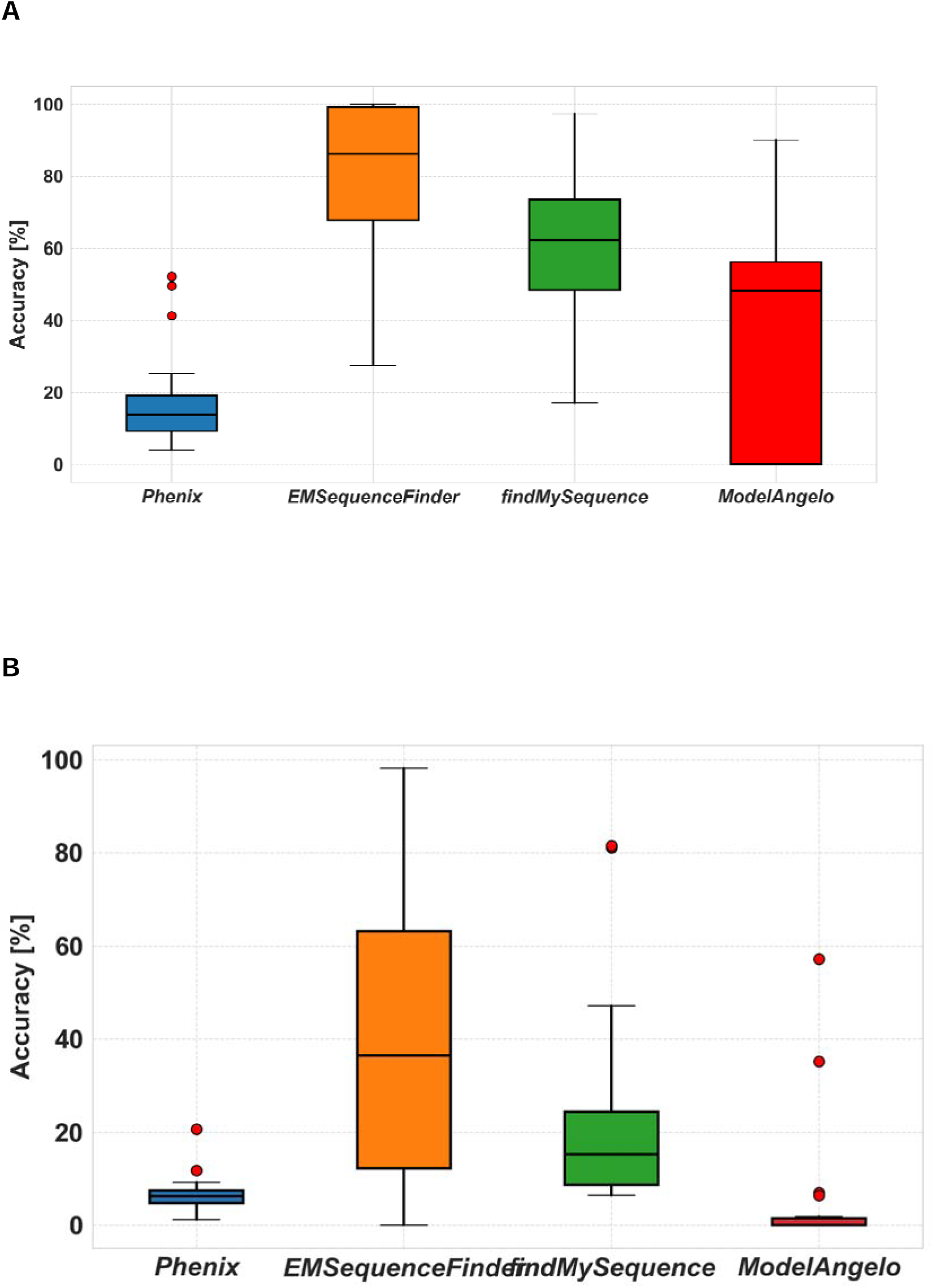
Comparison of *EMSequenceFinder* with state-of-the-art methods. Accuracies of assigning sequences to backbone traces for *EMSequenceFinder*, *Phenix*, ‘*findMySequence*’, and *ModelAngelo* for maps with resolution of (**A**) 4-4.5 L (Test set IIIa) and (B) 5-6 L (Test Set IIIb).

The observed accuracy differences can be attributed to the distinct goals and underlying assumptions of each method. *ModelAngelo* is a *de novo* model-building tool designed to construct complete atomic models directly from cryo-EM density maps, including challenging regions such as loops and poorly defined segments. In lower-resolution cases (e.g., Test Set IIIb), *ModelAngelo* often fails to generate complete models, resulting in fragmented outputs. When no assignable residues are identified, the apparent accuracy drops significantly, as meaningful comparisons to the reference structure are not possible. The *findMySequence* deep learning model was primarily trained on high-resolution data (≤L4.5LÅ), which likely contributes to its diminished accuracy on lower-resolution maps [23]. These methodological differences, including input requirements, training data, modeling strategies, and overall objectives, likely contribute to the observed accuracy differences.

### Impact of Backbone Perturbation and Scoring Prior on Sequence Assignment Accuracy

As described in **Methods**, backbone coordinates from deposited PDB structures were used to evaluate the performance of our scoring function. However, in practical applications, ground-truth models are not available. Fitting backbone traces into medium-resolution cryo-EM maps often results in alignment errors, ranging from small to severe. To assess the robustness of *EMSequenceFinde*r under such conditions, we evaluated its performance on selected maps in Test Set IIIa with perturbed backbone traces. For perturbation, we added random Gaussian-distributed coordinate displacements to the Cα atoms of each residue. Specifically, each Cartesian coordinate of the Cα atom was independently perturbed by a value drawn from a normal distribution with zero mean and a standard deviation of 0.5 Å or 1.0 Å. On average, accuracy decreased from 80.1% (unperturbed) to 61.1% with 0.5 Å deviation, and further to 42.6% with 1.0 Å deviation (Supplementary Table S2).

In addition to spatial inaccuracies, a more subtle challenge at intermediate resolutions is the ambiguity in determining the correct backbone directionality. To evaluate the ability of *EMSequenceFinder* to resolve these uncertainties, we simulated directional uncertainty by randomly reversing the residue order in 30% of the fragments, while keeping the remaining fragments unperturbed. The reversed and native fragments were then used together to recalculate the sequence assignment scores. Even with reversed fragments, *EMSequenceFinder* achieved an average accuracy of 45.8%.

Further, to evaluate the contribution of the secondary structure-based prior, we performed an ablation analysis by disabling the term from the scoring function. Accuracy was assessed across two resolution ranges: Test Set IIIa (4.0–4.5 Å) and Test Set IIIb (5.0–6.0 Å). In Test Set IIIa, average accuracy dropped from 77.6% (with prior) to 72.1% (without prior), while in Test Set IIIb, accuracy decreased from 39.5% to 37.4%. These results confirm that the secondary structure-based residue prior contributes meaningfully to the model’s robustness, especially at lower resolutions where side chain density is weak or ambiguous (Supplementary Fig. S3). To illustrate *EMSequenceFinder’s* accuracy across a range of resolutions, we visualized its predictions on two cryo-EM datasets: EMD-6526 (4.2 Å) and EMD-13360 (5.1 Å), taken from Test Sets IIIa and IIIb, respectively (Supplementary Fig. S4). STRIDE-based secondary structure fragments (helices and strands) from the reference PDB structures were compared to *EMSequenceFinder* predictions. In EMD-6526, a total of 54 secondary structure fragments were extracted. Among these, 26 fragments were assigned with perfect accuracy, 7 were partially matched, and 11 showed poor agreement. In contrast, for EMD-13360, despite the lower resolution, four well-resolved fragments were identified, all of which were correctly assigned.

While *EMSequenceFinder* performs relatively well across the tested resolution range, its accuracy depends on the accuracy of the backbone trace provided as input. Because the method evaluates sidechain density features relative to backbone atom positions, substantial deviations in backbone placement can lead to poor local density extraction, lowering the confidence of residue classification and sequence assignment. As *EMSequenceFinder* is not a backbone tracing tool but rather a threading method built on pre-fitted fragments, it assumes a reasonably accurate backbone model. This limitation highlights the importance of combining *EMSequenceFinder* with reliable backbone tracing approaches, particularly at lower resolutions.

### Comparison of SARS-CoV-2 Nsp2 sequence threadings by *EMSequenceFinder* and *findMySequence*

Finally, we illustrate *EMSequenceFinder* in a real world application by using it to annotate eight different cryo-EM maps of the SARS CoV-2 NSP2 protein reconstructed at resolutions between 3.7 and 7.0 L. A 3.9 Å reconstruction of Nsp2 was computed from images of ∼240,000 particles [44]. Further subclassification of this particle stack yielded a 3.7 Å map from ∼120,000 particles. To generate realistic lower resolution cryo-EM maps, smaller substacks from the ∼240,000 particle stack were used for refinement and reconstruction. Nsp2 maps were computed at 4.1 Å, 4.2 Å, 4.5 Å, 4.6 Å, 5.8 Å, and 7.0 Å resolutions reconstructed from 150,000, 100,000, 50,000, 20,000, 15,000, and 10,000 particles, respectively. All resolutions were estimated with a gold standard FSC protocol within a mask around the particle as implemented in cryoSPARC [45].

The backbone trace was obtained for the 3.7 Å map. This backbone trace was used to extract the sidechain densities as described in **Methods**. Because the deep learning methods are more accurate than *Phenix* (based on benchmarking with Test Set IIIa and Test Set IIIb), we only compared *findMySequence* and *EMSequenceFinder*. The accuracy of *EMSequenceFinder* deteriorates with map resolution, but we could still predict approximately 50% of the fragment sequences correctly at the map resolution of 6.0 L (**Fig. 7)**. While both methods become less accurate when the map resolution decreases below 4.5 L, the decrease in accuracy is worse for *findMySequence* than *EMSequenceFinder*, as already observed above.

**Figure 7.**
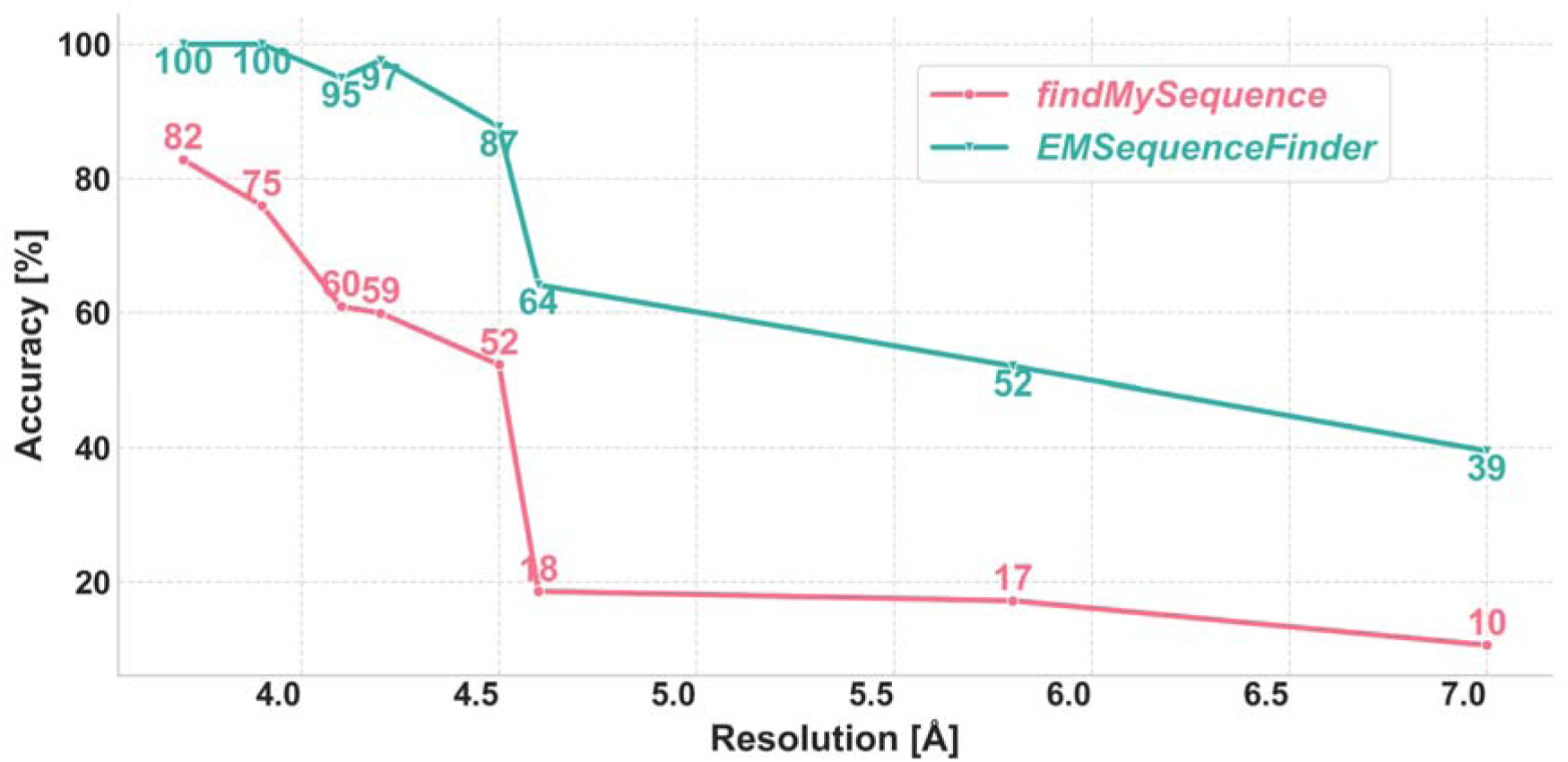
Accuracy of sequence prediction for different cryo-EM maps of the NSP2 protein as a function of map resolution.

## Conclusion

To facilitate building atomic protein structure models based on medium-resolution cryo-EM maps, we devised a deep learning method for assigning the amino acid residue sequence to the backbone fragments traced in an input cryo-EM map (*EMSequenceFinder*). *EMSequenceFinder* relies on a CNN trained on a large number of cryo-EM maps deposited in EMDB. *EMSequenceFinder* has a prediction accuracy of 41% for identifying individual amino acid residues, for maps at 3-8 L resolution. The prediction accuracy increases to 80% when predicting sequences of backbone traces, for maps at better than 5 Å resolution. Thus, we anticipate that *EMSequenceFinder* can be useful for computing models into maps at resolutions as low as ∼6 Å. The primary objective of this work is to evaluate whether the scoring function can accurately assign sequences when the backbone of a fragment is reasonably well-fitted. If the scoring function performs well under these conditions, dynamic programming or stochastic sampling methods (as mentioned in section **Databases for comparing *EMSequenceFinder* with state-of-the-art methods)** could be employed to simultaneously thread multiple fragments, potentially yielding a single optimal solution or an ensemble of plausible alternatives. In addition, the implementation of the *EMSequenceFinder* Bayesian scoring function in *IMP* allows it to be used in integrative modeling of multi-component protein structures, together with other information, such as models of some complex components and chemical crosslinks between them.

## Acknowledgments

Research was supported by NIH/NIGMS grants R01 GM083960 and P41 GM109824 (A.S.).

## Data Availability Statement

All datasets and code utilized in this study are publicly available at https://integrativemodeling.org/.

## References

1. Fukuda Y, Stapleton K, Kato T. Progress in spatial resolution of structural analysis by cryo-EM. Microscopy . 2023;72: 135–143.

2. Peplow M. Cryo-Electron Microscopy Reaches Resolution Milestone. ACS Cent Sci. 2020;6: 1274–1277.

3. Cowtan K. The Buccaneer software for automated model building. 1. Tracing protein chains. Acta Crystallogr D Biol Crystallogr. 2006;62: 1002–1011.

4. Terwilliger TC, Grosse-Kunstleve RW, Afonine PV, Moriarty NW, Zwart PH, Hung LW, et al. Iterative model building, structure refinement and density modification with the PHENIX AutoBuild wizard. Acta Crystallogr D Biol Crystallogr. 2008;64: 61–69.

5. Terashi G, Kihara D. De novo main-chain modeling for EM maps using MAINMAST. Nat Commun. 2018;9: 1618.

6. Liebschner D, Afonine PV, Baker ML, Bunkóczi G, Chen VB, Croll TI, et al. Macromolecular structure determination using X-rays, neutrons and electrons: recent developments in Phenix. Acta Crystallogr D Struct Biol. 2019;75: 861–877.

7. Alford RF, Leaver-Fay A, Jeliazkov JR, O’Meara MJ, DiMaio FP, Park H, et al. The Rosetta All-Atom Energy Function for Macromolecular Modeling and Design. J Chem Theory Comput. 2017;13: 3031–3048.

8. Pfab J, Phan NM, Si D. DeepTracer for fast de novo cryo-EM protein structure modeling and special studies on CoV-related complexes. Proc Natl Acad Sci U S A. 2021;118. doi:10.1073/pnas.2017525118

9. Jamali K, Käll L, Zhang R, Brown A, Kimanius D, Scheres SHW. Automated model building and protein identification in cryo-EM maps. bioRxiv. 2023. doi:10.1101/2023.05.16.541002

10. Zhong ED, Bepler T, Berger B, Davis JH. CryoDRGN: reconstruction of heterogeneous cryo-EM structures using neural networks. Nat Methods. 2021;18: 176–185.

11. Giri N, Cheng J. De novo atomic protein structure modeling for cryoEM density maps using 3D transformer and HMM. Nat Commun. 2024;15: 5511.

12. Terashi G, Wang X, Prasad D, Nakamura T, Kihara D. DeepMainmast: integrated protocol of protein structure modeling for cryo-EM with deep learning and structure prediction. Nat Methods. 2024;21: 122–131.

13. Barad BA, Echols N, Wang RY-R, Cheng Y, DiMaio F, Adams PD, et al. EMRinger: side chain-directed model and map validation for 3D cryo-electron microscopy. Nat Methods. 2015;12: 943–946.

14. Pintilie G, Zhang K, Su Z, Li S, Schmid MF, Chiu W. Measurement of atom resolvability in cryo-EM maps with Q-scores. Nat Methods. 2020;17: 328–334.

15. Nakamura T, Wang X, Terashi G, Kihara D. DAQ-Score Database: assessment of map-model compatibility for protein structure models from cryo-EM maps. Nat Methods. 2023;20: 775–776.

16. Casañal A, Shakeel S, Passmore LA. Interpretation of medium resolution cryoEM maps of multi-protein complexes. Curr Opin Struct Biol. 2019;58: 166–174.

17. Lawson CL, Patwardhan A, Baker ML, Hryc C, Garcia ES, Hudson BP, et al. EMDataBank unified data resource for 3DEM. Nucleic Acids Res. 2016;44: D396–403.

18. Terwilliger TC, Adams PD, Afonine PV, Sobolev OV. Cryo-EM map interpretation and protein model-building using iterative map segmentation. Protein Sci. 2020;29: 87–99.

19. Emsley P, Cowtan K. Coot: model-building tools for molecular graphics. Acta Crystallogr D Biol Crystallogr. 2004;60: 2126–2132.

20. Terwilliger TC. SOLVE and RESOLVE: automated structure solution and density modification. Methods Enzymol. 2003;374: 22–37.

21. LeCun Y, Bengio Y, Hinton G. Deep learning. Nature. 2015;521: 436–444.

22. Rawat W, Wang Z. Deep convolutional neural networks for image classification: A comprehensive review. Neural Comput. 2017;29: 2352–2449.

23. Chojnowski G, Simpkin AJ, Leonardo DA, Seifert-Davila W, Vivas-Ruiz DE, Keegan RM, et al. : a neural-network-based approach for identification of unknown proteins in X-ray crystallography and cryo-EM. IUCrJ. 2022;9: 86–97.

24. He J, Huang S-Y. Full-length de novo protein structure determination from cryo-EM maps using deep learning. Bioinformatics. 2021;37: 3480–3490.

25. Giri N, Roy RS, Cheng J. Deep learning for reconstructing protein structures from cryo-EM density maps: Recent advances and future directions. Curr Opin Struct Biol. 2023;79: 102536.

26. Avramov TK. Deep Learning for Validating Resolution and Detecting Secondary Structure Elements of Proteins in 3D Cryo-electron Microscopy Images. 2019.

27. Lagerstedt I, Moore WJ, Patwardhan A, Sanz-García E, Best C, Swedlow JR, et al. Web-based visualisation and analysis of 3D electron-microscopy data from EMDB and PDB. J Struct Biol. 2013;184: 173–181.

28. Webb B, Viswanath S, Bonomi M, Pellarin R, Greenberg CH, Saltzberg D, et al. Integrative structure modeling with the Integrative Modeling Platform: Integrative Structure Modeling with IMP. Protein Sci. 01/2018;27: 245–258.

29. Sali A. From integrative structural biology to cell biology. J Biol Chem. 2021;296: 100743.

30. Russel D, Lasker K, Webb B, Velázquez-Muriel J, Tjioe E, Schneidman-Duhovny D, et al. Putting the pieces together: integrative modeling platform software for structure determination of macromolecular assemblies. PLoS Biol. 2012;10: e1001244.

31. Baker ML, Ju T, Chiu W. Identification of secondary structure elements in intermediate-resolution density maps. Structure. 2007;15: 7–19.

32. Berman HM. The Protein Data Bank. Nucleic Acids Res. 2000;28: 235–242.

33. Kabsch W, Sander C. Dictionary of protein secondary structure: pattern recognition of hydrogen-bonded and geometrical features. Biopolymers. 1983;22: 2577–2637.

34. Chollet F. Keras GitHub. 2015. Available: https://github.com/fchollet/keras

35. Abadi M, Barham P, Chen J, Chen Z, Davis A, Dean J, et al. TensorFlow. Zenodo; 2023. doi:10.5281/ZENODO.4724125

36. Kingma DP, Ba J. Adam: A method for stochastic optimization. arXiv [cs.LG]. 2014. Available: http://arxiv.org/abs/1412.6980

37. Tee Y-Y, Cheng D, Chee C-S, Lin T, Shi Y, Gwee B-H. Unsupervised domain adaptation with histogram-gated Image Translation for delayered IC image analysis. arXiv [cs.CV]. 2022. Available: http://arxiv.org/abs/2209.13479

38. Heinig M, Frishman D. STRIDE: a web server for secondary structure assignment from known atomic coordinates of proteins. Nucleic Acids Res. 2004;32: W500–2.

39. Terwilliger TC, Sobolev OV, Afonine PV, Adams PD, Ho CM, Li X, et al. Protein identification from electron cryomicroscopy maps by automated model building and side-chain matching. Acta Crystallogr D Struct Biol. 2021;77: 457–462.

40. Pedregosa F, Varoquaux G, Gramfort A, Michel V, Thirion B, Grisel O, et al. Scikit-learn: Machine Learning in Python. arXiv [cs.LG]. 2012. pp. 2825–2830. Available: https://www.jmlr.org/papers/volume12/pedregosa11a/pedregosa11a.pdf?ref=https:/

41. Opitz J. A closer look at Classification evaluation metrics and a critical reflection of common evaluation practice. Trans Assoc Comput Linguist. 2024;12: 820–836.

42. He J, Li T, Huang S-Y. Improvement of cryo-EM maps by simultaneous local and non-local deep learning. Nat Commun. 2023;14: 3217.

43. Ho C-M, Li X, Lai M, Terwilliger TC, Beck JR, Wohlschlegel J, et al. Bottom-up structural proteomics: cryoEM of protein complexes enriched from the cellular milieu. Nat Methods. 2020;17: 79–85.

44. Gupta M, Azumaya CM, Moritz M, Pourmal S, Diallo A, Merz GE, et al. CryoEM and AI reveal a structure of SARS-CoV-2 Nsp2, a multifunctional protein involved in key host processes. bioRxiv; 2021. doi:10.1101/2021.05.10.443524

45. Punjani A, Rubinstein JL, Fleet DJ, Brubaker MA. cryoSPARC: algorithms for rapid unsupervised cryo-EM structure determination. Nat Methods. 2017;14: 290–296.

46. Bergstra J, Bengio Y. Random search for hyper-parameter optimization. J Mach Learn Res. 2012;13: 281–305.

