## Supplementary Information for "Recognizing amino acid sidechains in a medium resolution cryo-electron density map"

#### Hyperparameter Tuning

The architecture of the CNN model is described in the main text (**Fig. 2**). Here, we describe hyperparameter tuning. Several hyperparameters were evaluated on a subset of the training database to optimize the performance and computation time, while preserving the stability of learning. We evaluated the following parameters: learning rate, dropout rate, and the number of convolution blocks. A single parameter configuration was tested by randomly assigning a value within a set of bounds [[46]](https://paperpile.com/c/1nDGka/LUKRa). Each parameter configuration was applied to a representative random sample of 60% of the full dataset (this fraction was chosen as the best tradeoff between computation time and equilibration of model accuracy). For example, learning rates of 10^-1^, 10^-2^, …, 10^-6^ were tested, eventually choosing 0.00008 as the best balance of stability and convergence time. A batch size of 64 was chosen to allow for calculation on our 24GB Titan GPU hardware. We chose a maximum of 100 epochs for learning, with an early stopping criterion of 10 epochs without improvement of the loss function. A plot of improvement of the model accuracy and decrease of loss with respect to the number of epochs are shown in **Supplementary** **Figure S1**.

Additional Benchmarking Studies

1. Ablation Study: Removal of Secondary Structure-Based Prior

To assess the contribution of secondary structure-based priors (P(AA|SS)) to sequence assignment accuracy, we performed an ablation study where the priors were disabled. In this modified version of *EMSequenceFinder*, the prior probabilities based on secondary structure were omitted from the scoring function. Only the CNN-based sidechain density predictions were used to evaluate residue identities, without any weighting by expected secondary structure composition. The same backbone traces and sidechain density maps were used as inputs. The resulting sequence assignment accuracy was compared against the original model that included the priors (Supplementary Fig. S3).

2. Perturbation of Backbone Coordinates

To simulate inaccuracies in backbone tracing, Gaussian noise was added to the backbone atom coordinates (N, CA, C) of the reference PDB structures for selected EMDB maps from Test Set IIIa. Two levels of perturbation were applied independently, corresponding to 0.5 Å (moderate perturbation) and 1.0 Å standard deviation (severe perturbation), respectively. For each atom, random displacements were sampled from a normal distribution centered at zero with the specified standard deviation. After perturbation, *EMSequenceFinder* was run using the same input parameters, followed by evaluating the accuracy of sequence assignment (Supplementary Table S2).

3. Reversal of Sequence Direction

To evaluate the robustness of *EMSequenceFinder* against sequence directionality errors, we randomly reversed the residue order for 30% of the structure fragments generated for each fitted PDB structure for selected EMDB maps from Test Set IIIa, while keeping the atomic coordinates unchanged. For example, a fragment with residue sequence Ala–Val–Leu–Gly was reordered to Gly–Leu–Val–Ala, without altering the spatial positions of the atoms. The reversed fragments were then combined with the remaining unperturbed fragments to regenerate the input database files, while keeping the native density maps unmodified. Sequence assignment accuracy was recalculated on these perturbed sets and compared against the original unperturbed results. Fragments that became mismatched or unusable due to sequence reversal were treated as failed assignments. This approach allowed us to systematically test *EMSequenceFinder’s* sensitivity to fragment directionality errors (Supplementary Table S2).

### Supplementary Figures and Tables

**A**

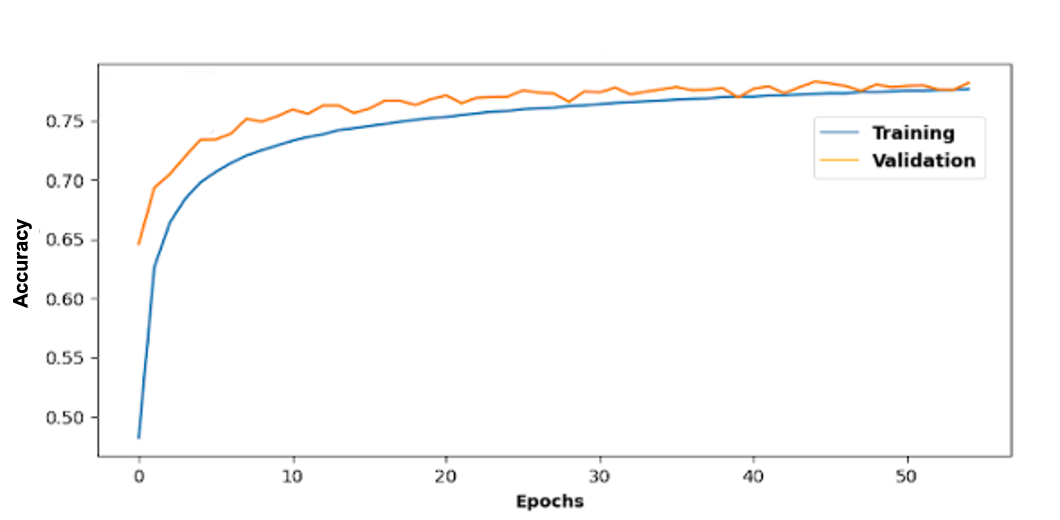

**B**

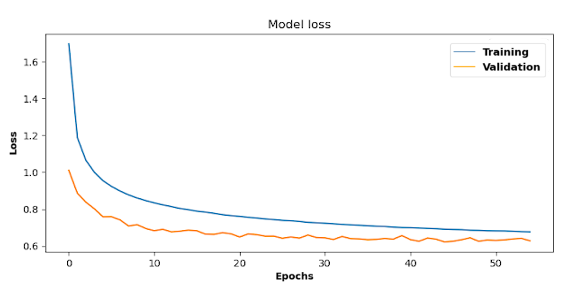

#### Supplemental Figure S1. Training of our final CNN model.

(**A**) Changes in prediction accuracy as a function of the number of epochs for our final CNN model. (**B**) Model loss as a function of the number of epochs for our final CNN model. Blue line, training and validation accuracy as a function of the number of epochs. Orange line, training and validation loss as a function of the number of epochs.

**A**

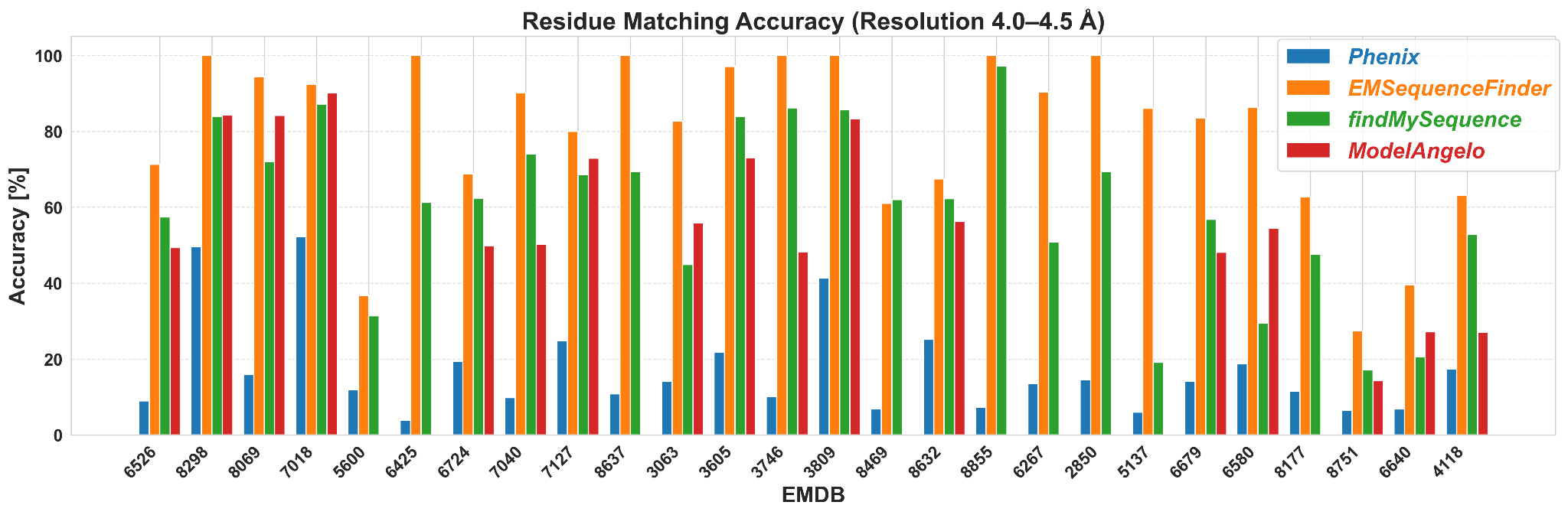

B

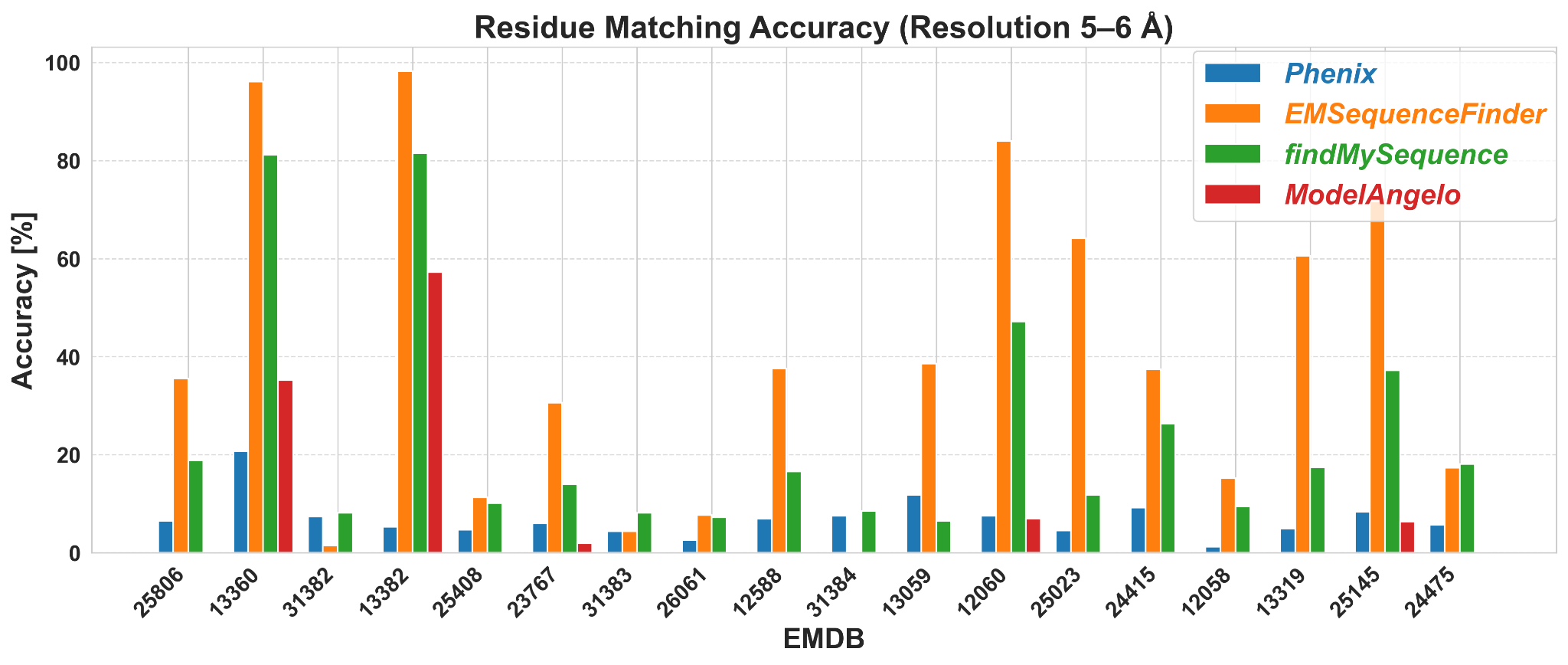

#### Supplemental Figure S2. Comparison of *EMSequenceFinder* with state-of-the-art methods.

Comparison of prediction accuracies for overall sequence threading by *EMSequenceFinder*, Phenix, *findMySequence* and *ModelAngelo* for maps with resolutions of (A) 4 - 4.5Å (Test Set IIIa) and (B) 5-6 Å (Test Set IIIb).

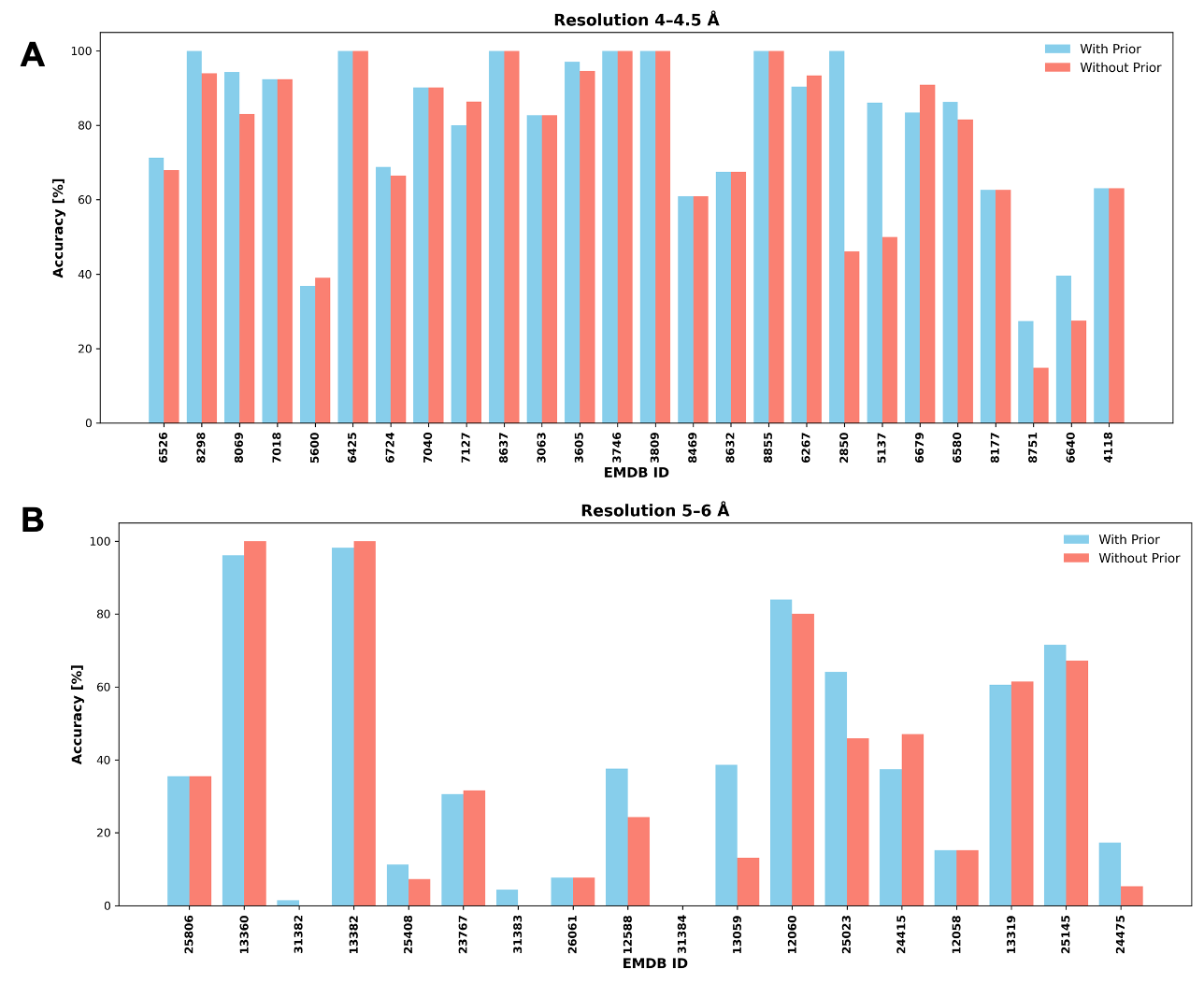

Supplementary Figure S3. Effect of secondary structure-based residue priors on sequence assignment accuracy. (A) 4 - 4.5Å (Test Set IIIa) and (B) 5-6 Å (Test Set IIIb).

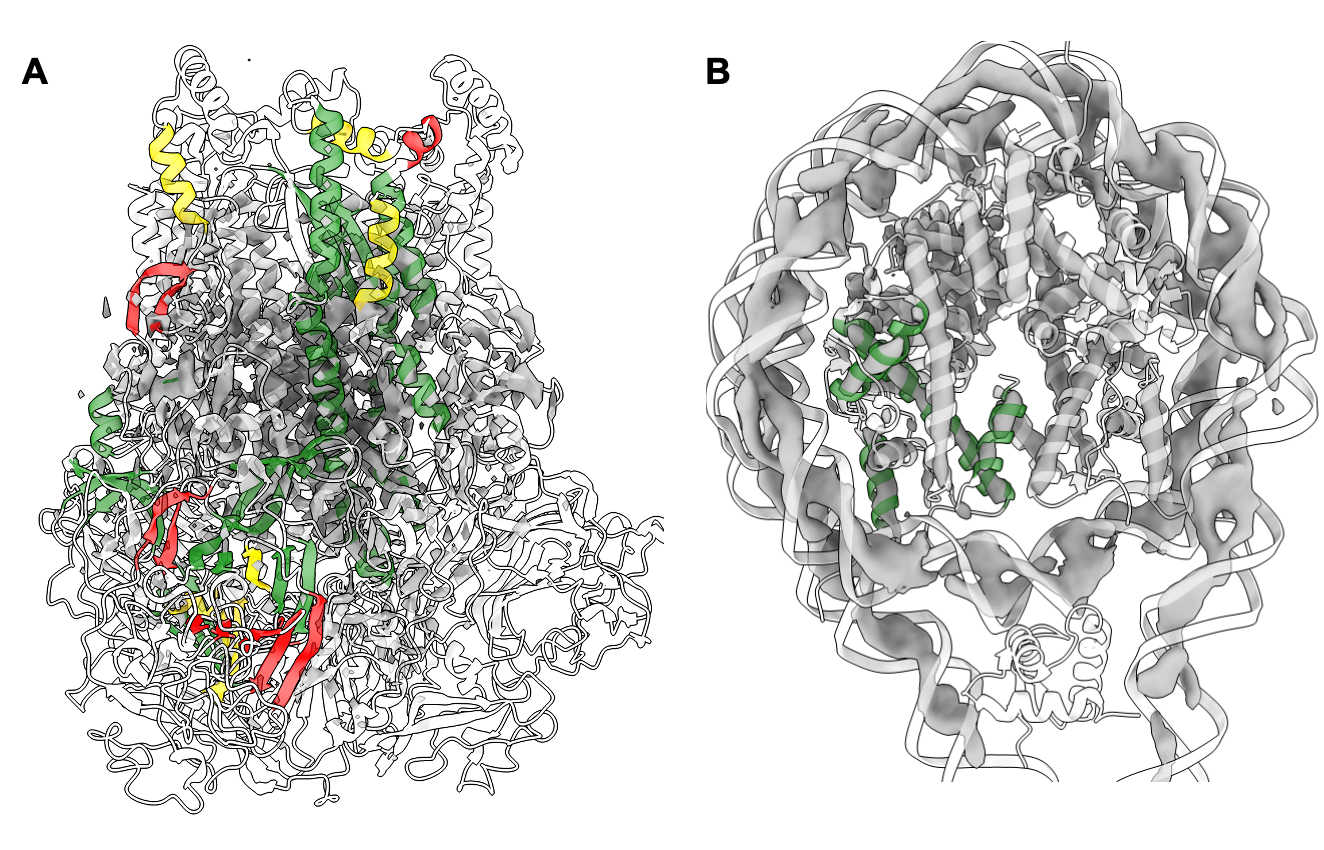

Supplementary Figure S4. Visualization of predicted and assigned fragments from EMSequenceFinder in fitted cryo-EM maps. (A) EMD-6526 (4.2 Å) and (B) EMD-13360 (5.1 Å). Correctly predicted fragments (rank = 0) are colored green, partial matches (0 < rank ≤ L/3) in yellow, and incorrect predictions/ambiguous backbone in red.

#### Supplemental Table S1. Test Sets IIIa and IIIb.

EMDB identifiers and resolutions of the cryo-EM maps in Test Sets IIIa (resolutions between 4 and 4.5 Å) and IIIb (resolutions between 5 and 6 Å).

| EMDB identifier | Resolution [Å] | Test Set |
| --- | --- | --- |
| 6526 | 4.0 | IIIa |
| 8298 | 4.0 |  |
| 8069 | 4.0 |  |
| 7018 | 4.1 |  |
| 5600 | 4.1 |  |
| 6425 | 4.1 |  |
| 6724 | 4.1 |  |
| 7040 | 4.1 |  |
| 7127 | 4.1 |  |
| 8637 | 4.1 |  |
| 3063 | 4.2 |  |
| 3605 | 4.2 |  |
| 3746 | 4.2 |  |
| 3809 | 4.2 |  |
| 8469 | 4.2 |  |
| 8632 | 4.2 |  |
| 8855 | 4.2 |  |
| 6267 | 4.2 |  |
| 2850 | 4.3 |  |
| 5137 | 4.3 |  |
| 6679 | 4.3 |  |
| 6580 | 4.4 |  |
| 8177 | 4.4 |  |
| 8751 | 4.4 |  |
| 6640 | 4.4 |  |
| 4118 | 4.5 |  |
| 25806 | 5.1 | IIIb |
| 13360 | 5.1 |  |
| 31382 | 5.1 |  |
| 13382 | 5.2 |  |
| 25408 | 5.2 |  |
| 23767 | 5.2 |  |
| 31383 | 5.3 |  |
| 26061 | 5.3 |  |
| 12588 | 5.3 |  |
| 31384 | 5.4 |  |
| 13059 | 5.4 |  |
| 12060 | 5.5 |  |
| 25023 | 5.5 |  |
| 24415 | 5.6 |  |
| 12058 | 5.7 |  |
| 13319 | 5.8 |  |
| 25145 | 5.9 |  |
| 24475 | 5.9 |  |

#### Supplemental Table S2.

Comparison of *EMSequenceFinder* accuracy on test set IIIa between original and perturbed backbone traces.

| EMDB Identifier | No Perturbation | Deviation_0.5 Å | Deviation_1.0 Å | Reversed Sequence |
| --- | --- | --- | --- | --- |
| 6526 | 71.3 | 63.373 | 72.34 | 54.581 |
| 8298 | 100.0 | 90.678 | 26.087 | 49.573 |
| 8069 | 94.4 | 85.443 | 57.471 | 70.96 |
| 7018 | 92.4 | 78.309 | 67.677 | 74.636 |
| 5600 | 36.8 | 34.975 | 9.615 | 36.164 |
| 6724 | 68.8 | 60.272 | 37.354 | 41.351 |
| 7127 | 80.0 | 91.101 | 35.115 | 65.517 |
| 8637 | 100.0 | 84.615 | 24.294 | 40.496 |
| 3605 | 97.1 | 82.692 | 100.0 | 73.301 |
| 3746 | 100.0 | 37.931 | 33.333 | 32.911 |
| 3809 | 100.0 | 80.556 | 70.833 | 32.609 |
| 8632 | 67.5 | 60.156 | 45.455 | 35.315 |
| 6267 | 90.4 | 93.413 | 93.413 | 76.647 |
| 2850 | 100.0 | 30.597 | 0.0 | 21.053 |
| 5137 | 86.1 | 32.824 | 0.0 | 31.481 |
| 6679 | 83.5 | 87.912 | 100.0 | 63.386 |
| 6580 | 86.3 | 43.277 | 33.735 | 48.246 |
| 8751 | 27.4 | 5.822 | 0.0 | 8.025 |
| 6640 | 39.6 | 17.384 | 2.825 | 13.447 |
